# R-spondins can potentiate WNT signaling without LGR receptors

**DOI:** 10.1101/198572

**Authors:** Andres M. Lebensohn, Rajat Rohatgi

## Abstract

The WNT signaling pathway regulates patterning and morphogenesis during embryonic development and promotes tissue renewal and regeneration in adults. Some WNT responses in vertebrates depend on a second signal provided by the R-spondin family of four secreted proteins (RSPO1-4) that drive the renewal of stem cells in many tissues. RSPOs markedly amplify target cell sensitivity to WNT ligands by neutralizing two transmembrane E3 ligases, ZNRF3 and RNF43, which reduce cell-surface levels of WNT receptors. Chromosomal translocations that increase RSPO expression or that inactivate ZNRF3/RNF43 can drive human cancers. RSPOs contain tandem furin-like repeats (FU1 and FU2), a thrombospondin type I (TSP) domain, and a basic region (BR). RSPOs simultaneously engage ZNRF3/RNF43 through their FU1 domain and one of three leucine-rich repeat-containing G-protein coupled receptors (LGR4-6) through their FU2 domain, triggering the clearance of ZNRF3/RNF43 and the consequent rise in WNT receptor levels. LGRs are selectively expressed in various tissue stem cells and are considered the primary high-affinity receptors for RSPOs. Using purified mutant and chimeric RSPOs and cell lines lacking various receptors, we show that RSPO2 and RSPO3, but not RSPO1 and RSPO4, can potentiate WNT/β-catenin signaling in the absence of all three LGRs. The ZNRF3/RNF43-interacting FU1 domain was necessary for LGR-independent signaling, while the LGR-interacting FU2 domain was dispensable. The FU1 domain of RSPO3 was also sufficient to confer LGR-independence when transplanted to RSPO1, demonstrating that its interaction with ZNRF3/RNF43 dictates LGR-independent signaling. The enigmatic TSP/BR domains of RSPOs and their interaction with heparan sulfate proteoglycans (HSPGs), previously considered dispensable for WNT/β-catenin signaling, became essential in the absence of LGRs. These results define two alternative modes of RSPO-mediated signaling that share a common dependence on ZNRF3/RNF43, but differ in their use of either LGRs or HSPGs, with implications for understanding their mechanism of action, biological functions and evolutionary origins.

## Text

The WNT signaling pathway regulates patterning and morphogenesis during embryonic development and promotes tissue renewal and regeneration in adults^1,2^. Some WNT responses in vertebrates depend on a second signal provided by the R-spondin family of four secreted proteins (RSPO1-4) that drive the renewal of stem cells in many tissues^3,4^. RSPOs markedly amplify target cell sensitivity to WNT ligands by neutralizing two transmembrane E3 ligases, ZNRF3 and RNF43, which reduce cell-surface levels of WNT receptors^5,6^. Chromosomal translocations that increase RSPO expression or that inactivate ZNRF3/RNF43 can drive human cancers^7^. RSPOs contain tandem furin-like repeats (FU1 and FU2), a thrombospondin type I (TSP) domain, and a basic region (BR). RSPOs simultaneously engage ZNRF3/RNF43 through their FU1 domain and one of three leucine-rich repeat-containing G-protein coupled receptors (LGR46) through their FU2 domain^8–12^, triggering the clearance of ZNRF3/RNF43 and the consequent rise in WNT receptor levels. LGRs are selectively expressed in various tissue stem cells and are considered the primary high-affinity receptors for RSPOs^13–15^. Using purified mutant and chimeric RSPOs and cell lines lacking various receptors, we show that RSPO2 and RSPO3, but not RSPO1 and RSPO4, can potentiate WNT/β-catenin signaling in the absence of all three LGRs. The ZNRF3/RNF43-interacting FU1 domain was necessary for LGR-independent signaling, while the LGR-interacting FU2 domain was dispensable. The FU1 domain of RSPO3 was also sufficient to confer LGR-independence when transplanted to RSPO1, demonstrating that its interaction with ZNRF3/RNF43 dictates LGR-independent signaling. The enigmatic TSP/BR domains of RSPOs and their interaction with heparan sulfate proteoglycans (HSPGs), previously considered dispensable for WNT/β-catenin signaling^16,17^, became essential in the absence of LGRs. These results define two alternative modes of RSPO-mediated signaling that share a common dependence on ZNRF3/RNF43, but differ in their use of either LGRs or HSPGs, with implications for understanding their mechanism of action, biological functions and evolutionary origins.

In previous work^18^, we generated and thoroughly characterized a haploid human cell line (HAP1-7TGP) that harbors a fluorescent transcriptional reporter for WNT/β-catenin signaling. Both the fluorescence of this synthetic reporter and the transcription of endogenous WNT target genes in HAP1-7TGP cells can be activated by WNT ligands, and these WNT responses can be strongly potentiated by RSPOs^18^. HAP1-7TGP cells do not secrete WNT ligands and thus their response to RSPOs requires the co-administration of a low concentration of WNT. A comprehensive set of unbiased genetic screens in HAP1-7TGP identified most of the known components required for a signaling response to RSPO and WNT ligands, establishing this cell line as a valid and genetically tractable system for the study of this pathway^18^.

We made the serendipitous and unexpected observation that RSPO3 could potently enhance WNT reporter fluorescence driven by a low concentration of WNT3A in two independently derived HAP1-7TGP clonal cell lines carrying loss of function mutations in *LGR4* (LGR4^KO^ cells; see Methods and Supplementary Data File 1) (Fig. 1a). In contrast, RSPO1 was inactive in LGR4^KO^ cells. RSPO1 and RSPO3 had equivalent activity in wild-type (WT) HAP1-7TGP cells, demonstrating that both ligands were active, and responses in both WT and LGR4^KO^ cells depended on the presence of WNT3A (Fig. 1a).

**Figure 1.**
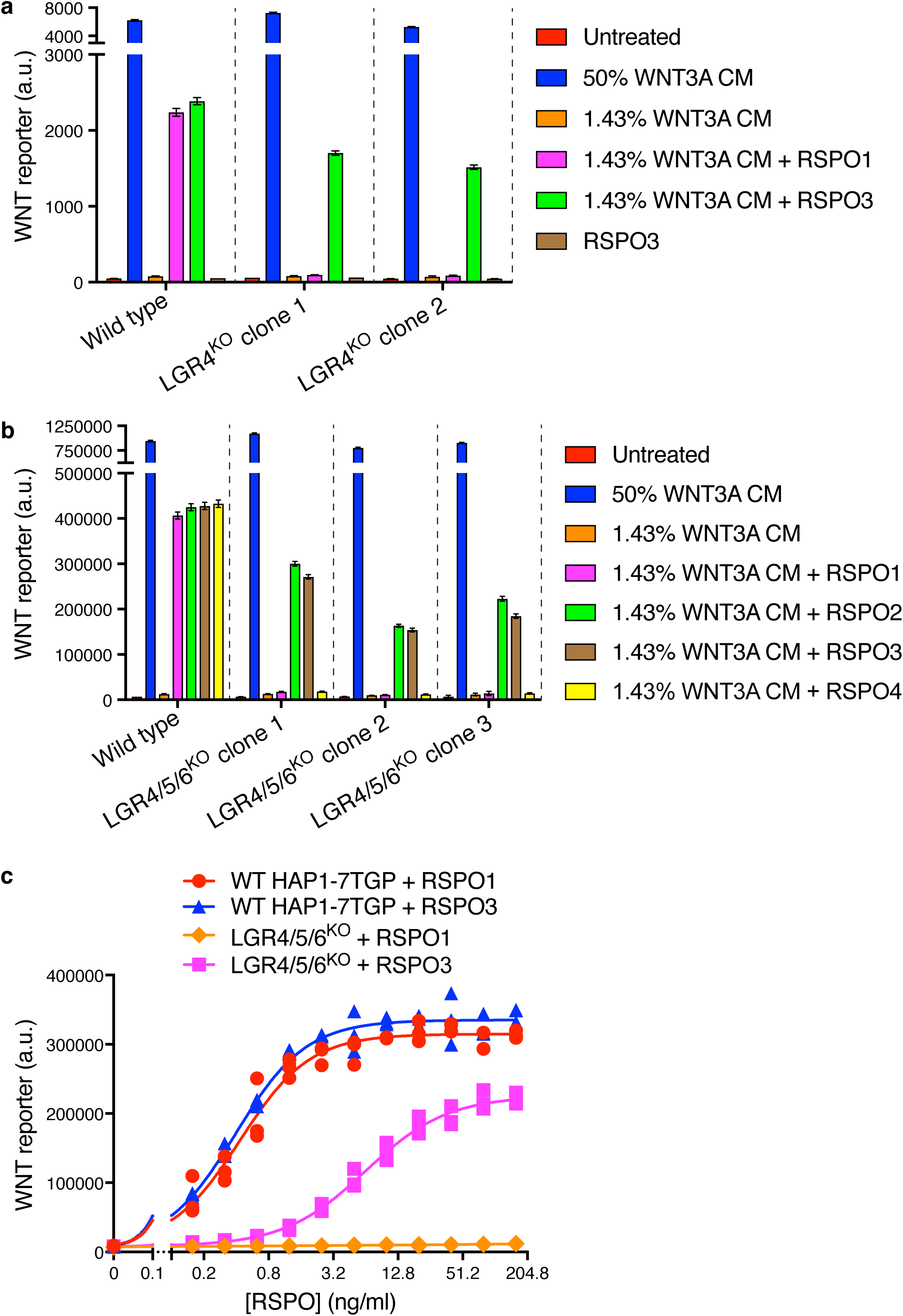
RSPO2 and RSPO3 can potentiate WNT signaling in the absence of LGR4, LGR5 and LGR6. a. WNT reporter fluorescence (median +/- standard error of the median (SEM) from 10,000 cells) for WT HAP1-7TGP and two LGR4^KO^ clonal cell lines following treatment with the indicated combinations of WNT3A conditioned media (CM) and untagged, recombinant RSPO1 or RSPO3 (both at 20 ng/ml). All cell lines responded similarly to a saturating dose of WNT3A, demonstrating an intact downstream signaling response. b. WNT reporter fluorescence (median +/- SEM from 10,000 cells) for WT HAP1-7TGP and three LGR4/5/6^KO^ clonal cell lines treated with the indicated combinations of WNT3A CM and various RSPOs. RSPO1, RSPO2 and RSPO3 were used at 40 ng/ml and RSPO4 at 400 ng/ml, concentrations that produced equivalent responses in WT cells. c. Dose-response curves for RSPO1 and RSPO3 in WT HAP1-7TGP and LGR4/5/6^KO^ cells in the presence of 1.43% WNT3A CM. Each symbol represents the median WNT reporter fluorescence from 5,000 cells in a single well, and three independently treated wells were measured for each RSPO concentration. The curves were fitted as described in Methods.

While *LGR4* is the only RSPO receptor expressed in HAP1 cells (Extended Data Table 1), we excluded the possibility of compensatory up-regulation of *LGR5* or *LGR6* by simultaneously disrupting both genes in LGR4^KO^ cells, generating multiple independent clonal cell lines lacking all three RSPO receptors (hereafter called LGR4/5/6^KO^ cells; Supplementary Data File 1). LGR4/5/6^KO^ cells retained an intact WNT signaling cascade, responding normally to a saturating dose of WNT3A (Fig. 1b). All four RSPOs strongly potentiated WNT signaling in WT cells, establishing ligand activity. However, RSPO1 and RSPO4 were completely inactive in LGR4/5/6^KO^ cells, even at concentrations that induced maximum WNT reporter induction in WT cells, whereas RSPO2 and RSPO3 strongly potentiated signaling driven by low concentrations of WNT3A even in the absence of all three LGR receptors (Fig. 1b). Therefore, RSPO2 and RSPO3 possess a unique quality absent in RSPO1 and RSPO4 that enables them to potentiate WNT responses without LGR receptors.

Dose-response analysis revealed that RSPO1 and RSPO3 enhanced WNT signaling in WT cells with nearly identical pharmacodynamics—both the efficacy (maximum reporter activity) and the potency (measured by the EC50, defined as the RSPO concentration that induced half-maximum reporter activity) were similar for both ligands (Fig. 1c). In LGR4/5/6^KO^ cells, RSPO1 had no detectable activity at concentrations up to 160 ng/ml, 400-fold higher than its EC50 (0.4 ng/ml) in WT cells. While RSPO3 potentiated WNT signaling in LGR4/5/6^KO^ cells, its efficacy was reduced by 33% and its EC50 was increased by 16-fold compared to WT cells (6.4 ng/ml vs. 0.4 ng/ml; Fig. 1c). The distinct RSPO3 pharmacodynamics in the two cell types suggested that the reception of RSPO3 was mediated by different mechanisms in the presence and absence of LGR receptors.

We sought to determine which domains of RSPO3 were required for LGR-independent signaling using a ligand mutagenesis strategy (Fig. 2a). Our experimental strategy leveraged a comparison between RSPO1 and RSPO3, since the former depended strictly on LGR receptors while the latter could signal in their absence. Unless otherwise noted, each WT and mutant RSPO ligand described hereafter was produced as a fusion protein carrying an N-terminal hemagglutinin (HA) tag and a tandem C-terminal tag composed of an immunoglobulin fragment crystallizable (Fc) domain followed by a 1D4 epitope tag^19^ used for immuno-affinity purification (see Extended Data Fig. 1a, b, and Methods for a description of ligand purification and characterization). Importantly, the pharmacodynamics of the tagged RSPO proteins were similar to those of their untagged counterparts in both WT and LGR4/5/6^KO^ cells (Extended data Fig. 1c, d).

**Figure 2.**
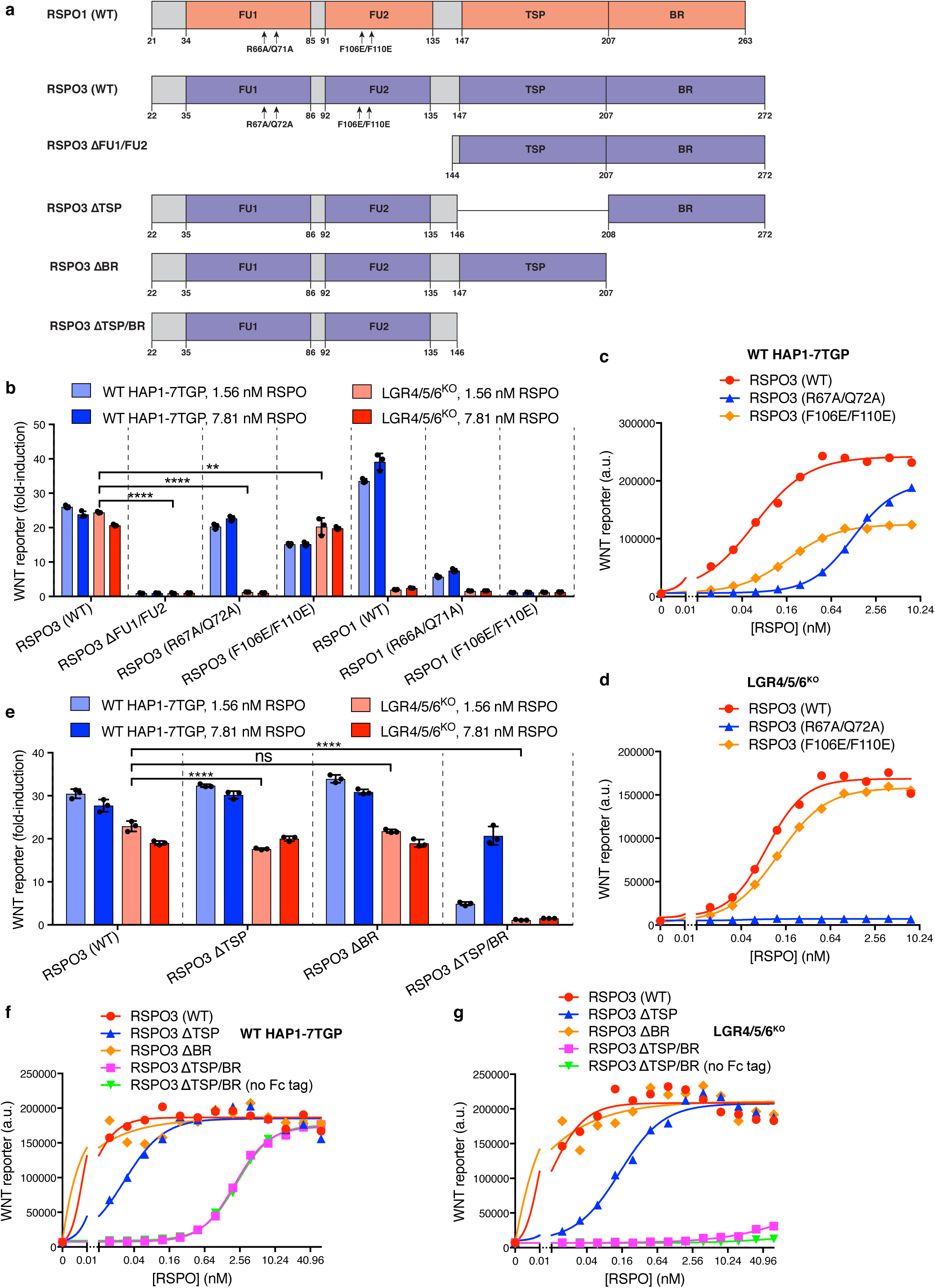
Domains of RSPO3 required for LGR-independent signaling. a. Schematic representation of human WT and mutant RSPO1 (pink) and RSPO3 (light blue) proteins produced and purified as described in Methods and Extended Data Figure 1a. The N-terminal HA and the C-terminal Fc and 1D4 tags present in all constructs are not shown. Amino acid numbers for human RSPO1 and RSPO3 (UniProt accession number Q2MKA7 and Q9BXY4, respectively) are indicated below, and arrows show the sites of mutations in the FU1 and FU2 domains. Polypeptide lengths are drawn to scale. b and e. Fold-induction in WNT reporter fluorescence over 1.43% WNT3A CM alone (bars and error bars indicate the average +/- SD from triplicate wells; circles indicate the fold-induction for individual wells) in WT HAP1-7TGP (light blue and blue bars) and LGR4/5/6^KO^ (pink and red bars) cells treated with two concentrations of purified RSPO proteins. Significance was determined as described in Methods. c, d, f, g. Dose-response curves for the indicated purified RSPO proteins in WT HAP1-7TGP (c,f) and LGR4/5/6^KO^ (d, g) cells in the presence of 1.43% WNT3A CM. Each symbol represents the median WNT reporter fluorescence from 5,000 cells. In f and g, RSPO3 ΔTSP/BR was tested with and without the dimerizing Fc tag.

Previous studies have shown that the FU1 and FU2 domains in all RSPOs, which bind to ZNRF3/RNF43 and LGRs, respectively, are both necessary and sufficient to potentiate WNT responses, while the TSP and BR domains are dispensable^16^. Simultaneous deletion of the FU1 and FU2 domains of RSPO3 abolished signaling in both WT and LGR4/5/6^KO^ cells (Fig. 2b). Point mutations in the FU1 domain (R67A/Q72A; Fig. 2a) known to weaken the interaction between RSPOs and ZNRF3/RNF43^20^ entirely abolished RSPO3 signaling in LGR4/5/6^KO^ cells (Fig. 2b, d) and substantially reduced (but did not abolish) RSPO3 signaling in WT cells (EC50 increased by 21-fold; Fig. 2b, c). Thus, the reduction in the affinity between RSPO3 and ZNRF3/RNF43 caused by the FU1 R67A/Q72A mutation impaired LGR-independent signaling to a much greater extent than LGR-dependent signaling. Indeed, the equivalent R66A/Q71A mutation in RSPO1 (Fig. 2a), which only signals in an LGR-dependent manner, also impaired but did not completely abolish signaling in WT cells (Fig. 2b).

Point mutations in the FU2 domain (F106E/F110E; Fig. 2a) of RSPO3 that weaken interactions with LGR receptors^20^ had little impact on RSPO3 signaling in LGR4/5/6^KO^ cells, consistent with the lack of LGRs in these cells (Fig. 2b, d). In WT cells, the F106E/F110E mutation in RSPO3 did not prevent signaling, but reduced the efficacy by 48% and increased the EC50 by 2.9-fold (Fig. 2c). Thus, RSPO3 signaling in WT cells includes contributions from both LGR-dependent and independent pathways. In contrast, the F106E/F110E mutation in RSPO1 abolished signaling in WT cells, demonstrating that signaling by RSPO1 depends entirely on its interaction with LGR receptors (Fig. 2b).

The C-terminal TSP and BR domains of RSPOs (denoted TSP/BR when discussed together) are considered dispensable for LGR-mediated signaling^15^. When we deleted these domains individually in RSPO3, there were only minor effects on signaling in WT cells (Fig. 2e, f). Deletion of both domains in RSPO3 increased the EC50 in WT cells by 333-fold, but did not change the efficacy (Fig. 2f). The signaling properties of RSPO3 lacking the TSP/BR domains were unchanged when the dimerizing Fc tag was removed (Fig. 2f). Therefore, while the TSP/BR domains are not essential for signaling in WT cells, consistent with prior work, their loss substantially reduces the apparent potency of RSPO3. In contrast, RSPO3 lacking the TSP/BR domains could not potentiate WNT responses in LGR4/5/6^KO^ cells (Fig.2e, g).

These mutagenesis experiments demonstrated that the FU1 and TSP/BR domains of RSPO3 are required for its ability to potentiate WNT responses in the absence of LGR receptors, while the FU2 domain is dispensable. These domain requirements are distinct from those for LGR-mediated signaling by RSPO1, which depends on the FU1 and FU2, but not on the TSP/BR domains. In WT cells, RSPO3 signaling proceeded through both LGR-dependent and independent mechanisms because disruption of the FU2 or the TSP/BR domains partially impaired but did not abolish signaling (Fig. 2c, f). The ZNRF3/RNF43-interacting FU1 domain is essential for signaling both in the presence and absence of LGR receptors.

To identify the region of RSPO3 that confers the property of LGR-independent signaling, we constructed a series of chimeric ligands, combining regions of RSPO1 and RSPO3 (Fig. 3a). Remarkably, replacing the FU1 domain of RSPO1 with the FU1 domain of RSPO3 enabled RSPO1 to potentiate WNT signaling in LGR4/5/6^KO^ cells (Fig. 3b, d). Conversely, replacing the FU1 domain of RSPO3 with that of RSPO1 drastically reduced the signaling capacity of RSPO3 in LGR4/5/6^KO^ cells (Fig. 3b, d). In important control experiments, all chimeric ligands showed equivalent activity in WT cells, establishing ligand integrity (Fig. 3b, c). Thus, a difference in the interaction between ZNRF3/RNF43 and the FU1 domains of RSPO1 and RSPO3 is the crucial determinant of LGR-independent signaling. Of note, the affinities of the FU1-FU2 fragment of RSPO2 (25 nM) and RSPO3 (60 nM) for ZNRF3 have been reported to be much higher than those of RSPO1 (6.8 μM) and RSPO4 (300 μM)^12^. Indeed, these affinities correlate with the capacity of RSPO2 and RSPO3, but not RSPO1 or RSPO4, to promote LGR-independent signaling (Fig. 1b). While the TSP/BR domains of RSPO3 were required for LGR-independent signaling, they were not sufficient because replacement of the TSP/BR domains of RSPO1 with those of RSPO3 did not confer the capacity to signal in LGR4/5/6^KO^ cells (Fig. 3e). In fact, the TSP/BR domains of RSPO1 and RSPO3 seemed interchangeable for signaling activity in both WT and LGR4/5/6^KO^ cells (Fig. 3e).

**Figure 3.**
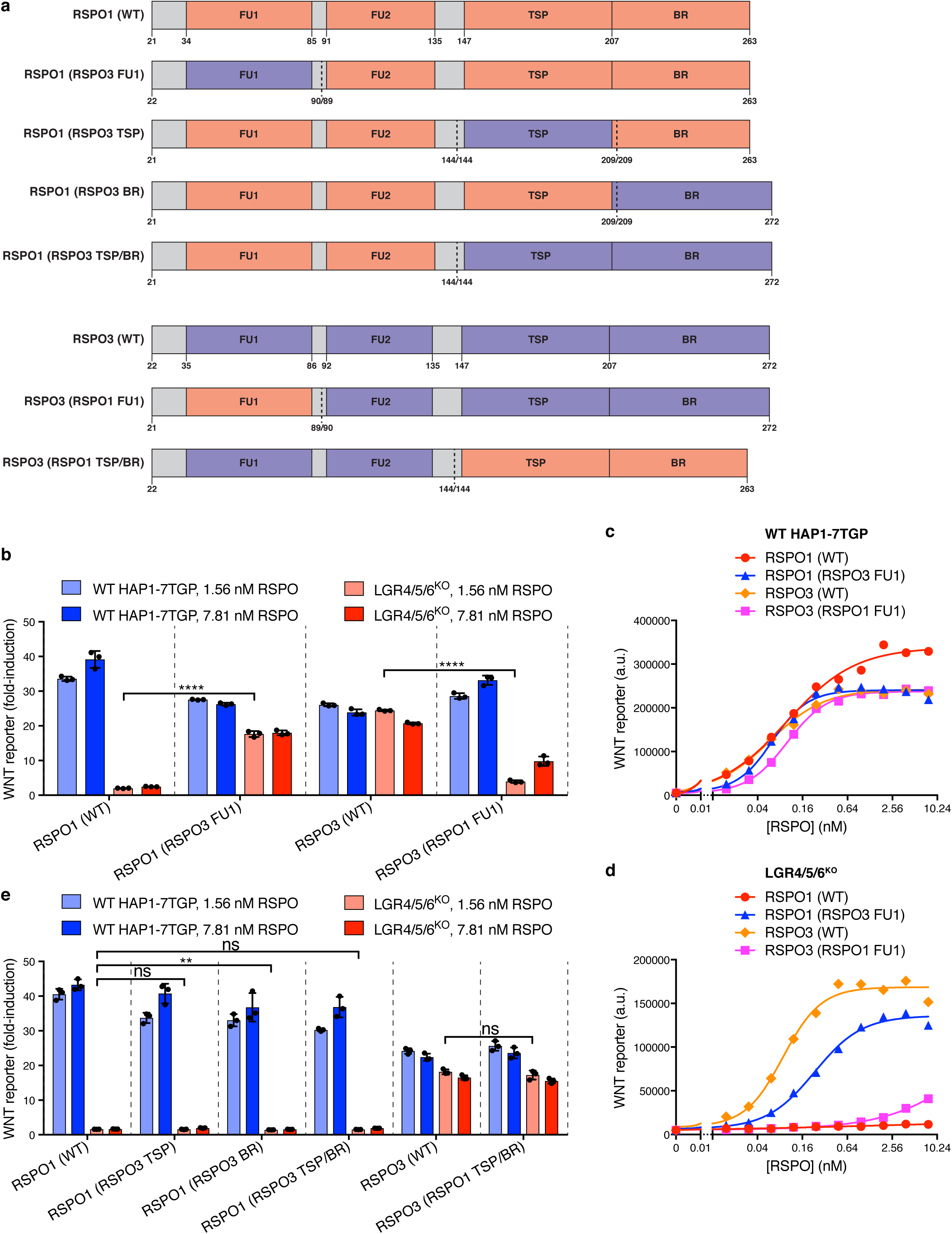
The FU1 domain of RSPO3 is sufficient to confer LGR-independent signaling when transplanted to RSPO1. a. Schematic representation of human WT and chimeric RSPO1 (pink) and RSPO3 (light blue) proteins, depicted as in Fig. 2a. Vertical dotted lines indicate the sites at which the swaps were made. Each swap was made at a conserved residue, whose number on the left and right of the slash corresponds to the protein depicted on the left and right of the dotted line, respectively. b and e. Fold-induction in WNT reporter fluorescence over 1.43% WNT3 CM alone (bars and error bars indicate the average +/- SD from triplicate wells; circles indicate the fold-induction for individual wells) in WT HAP1-7TGP (light blue and blue bars) and LGR4/5/6^KO^ (pink and red bars) cells treated with two concentrations of purified RSPO proteins. Significance was determined as described in Methods. c and d. Dose-response curves for the indicated purified RSPO proteins in WT HAP1-7TGP (c) and LGR4/5/6^KO^ (d) cells in the presence of 1.43% WNT3A CM. Each symbol represents the median WNT reporter fluorescence from 5,000 cells.

These results suggested that the WNT-potentiating activity of RSPO3 in the absence of the LGRs depends on its interaction with ZNRF3/RNF43 through the FU1 domain and an additional interaction with an alternative co-receptor through the TSP/BR domains. We considered the previous observation that the TSP/BR domains of RSPOs can bind to heparin^21^. Addition of heparin to the culture medium completely blocked signaling by RSPO3 in LGR4/5/6^KO^ cells, but had only a partial inhibitory effect on WT cells, in which RSPO3 can also signal through LGRs (Fig. 4a).

**Figure 4.**
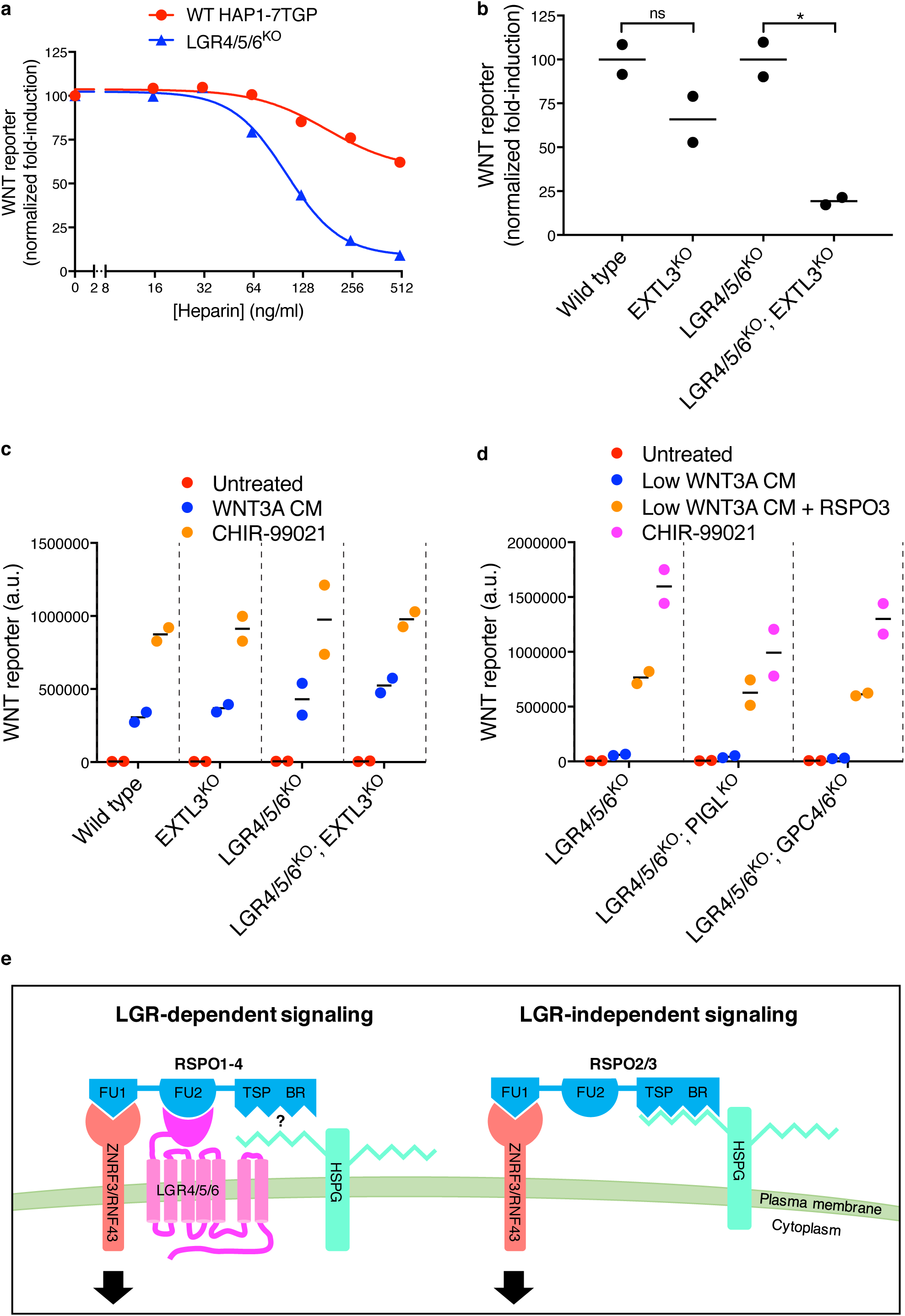
LGR-independent signaling by RSPO3 requires heparan sulfate proteoglycans. a. WNT reporter induction (calculated from the median WNT reporter fluorescence from 5,000 cells) in WT HAP1-7TGP and LGR4/5/6^KO^ cells stimulated with 1.43% WNT3A CM, 2 nM untagged RSPO3 and the indicated concentrations of heparin. The fold-induction over 1.43% WNT3A CM alone in the absence of heparin was normalized to 100% b. WNT reporter induction (calculated from the average WNT reporter fluorescence of triplicate wells) in the indicated cell lines following treatment with 2.78% WNT3A CM and 20 ng/ml untagged RSPO3. The fold-induction was normalized to the average fold-induction for WT (left two genotypes) or for LGR4/5/6^KO^ (right two genotypes) cells. Each circle represents a unique clonal cell line (determined by genotyping, Supplementary Data File 1) and the average of data from two independent clonal cell lines for each genotype is indicated by a horizontal line. Significance was determined as described in Methods. c.WNT reporter fluorescence (average +/- SD from triplicate wells) for the same clonal cell lines depicted in b. Where indicated, cells were treated with a sub-saturating concentration (11.1%) of WNT3A CM or with 10 µM of the GSK3 inhibitor CHIR-99021. d. WNT reporter fluorescence (average +/- SD from duplicate wells) following treatment with a low concentration of WNT3A CM (2.78% for LGR4/5/6^KO^ cells, or 11.1% for LGR4/5/6^KO^; PIGL^KO^ and LGR4/5/6^KO^; GPC4/6^KO^ cells) alone or in combination with 20 ng/ml RSPO3, or with 10 μM of the GSK3 inhibitor CHIR-99021. Since depletion of PIGL or of GPC4 and GPC6 reduces signaling at low WNT concentrations^18^, different WNT3A CM concentrations were used to achieve comparable signaling responses to WNT3A alone in all cell lines so that potentiation by the further addition of RSPO3 could be directly compared. Each circle represents a unique clonal cell line, and the average of data from two independent clonal cell lines for each genotype is indicated by a horizontal line. e. Proposed models for LGR-dependent and LGR-independent signaling by RSPOs. See text for details.

The TSP/BR domains of RSPOs can mediate interactions with the two major families of cell-surface heparan sulfate proteoglycans (HSPGs), the transmembrane syndecans and the glycophosphatidylinositol (GPI)-linked glypicans^17^. In humans, both protein families are encoded by multiple, partially redundant genes: four syndecan genes (*SDC1-4*) and six glypican genes (*GPC1-6*)^22^, all of which are expressed in HAP1 cells (Extended Data Table 1), making their genetic analysis challenging. Since all syndecans and glypicans must be post-translationally modified with heparan sulfate chains for receptor function, we disrupted *EXTL3*, a gene encoding a glycosyltransferase that is specifically required for HSPG biosynthesis, but dispensable for the synthesis of other glycosaminoglycans and proteoglycans^23^. The loss of EXTL3 in LGR4/5/6^KO^ cells led to an 81% reduction in RSPO3-mediated potentiation of WNT signaling (Fig. 4b). In contrast, the loss of EXTL3 in WT cells only reduced signaling by 34%, likely because RSPO3 can also signal through LGR receptors in WT cells. In an important control, the loss of EXTL3 did not affect signaling induced by addition of WNT3A alone or by inhibition of the β-catenin destruction complex kinase GSK3 in either WT or LGR4/5/6^KO^ cells (Fig. 4c).

To distinguish between syndecans and glypicans, we took advantage of the fact that only glypicans are anchored to the cell surface by a GPI linkage. Disrupting *PIGL*, a gene required for GPI-anchor biosynthesis, or disrupting both *GPC4* and *GPC6* (the two glypican genes identified in our previous haploid genetic screens^18^) in LGR4/5/6^KO^ cells did not impair LGR-independent potentiation of WNT signaling by RSPO3 (Fig. 4d).

These results suggest that the interaction of the TSP/BR domains of RSPO3 with cell surface HSPGs, possibly syndecans, provides an alternative mechanism that neutralizes ZNRF3/RNF43 in the absence of LGR receptors (Fig. 4e). Cell-surface HSPGs are known to mediate the efficient endocytosis of multiple cargoes^24^. Hence, we speculate that the simultaneous interaction of RSPO3 with ZNRF3/RNF43 through its FU1 domain and cell surface HSPGs through its TSP/BR domains provides an LGR-independent route for the endocytosis and clearance of ZNRF3/RNF43 from the cell surface (Fig. 4e), and the consequent rise in WNT receptor levels.

Our work shows that RSPOs can potentiate WNT signals in the absence of LGR receptors, expression of which has been hitherto considered the hallmark of RSPO-responsive cells. Future work will define the developmental, regenerative, and oncogenic contexts in which this LGR-independent mode of signaling is used to amplify target cell responses to WNT ligands. The mutant and chimeric RSPO ligands we described should allow the selective modulation of these alternate modes of signaling to dissect their biological roles.

## Methods

The following materials and methods relevant to this manuscript have been described previously ^18^: cell lines and growth conditions, preparation of WNT3A conditioned medium (CM) and construction of the HAP1-7TGP WNT reporter haploid cell line.

### Plasmids

pCX-Tev-Fc (unpublished) was a gift from Henry Ho (University of California Davis, Davis, CA). pHLsec-HA-Avi-1D4 (unpublished, derived from pHLSec ^25^ by incorporating a C-terminal HA tag following the signal sequence, and an N-terminal Gly/Ser linker, AviTag biotinylation sequence and 1D4 tag^19^) was a gift from Christian Siebold (University of Oxford, Oxford, United Kingdom). RSPO1-GFP^26^ was a gift from Feng Cong (Developmental and Molecular Pathways, Novartis Institutes for Biomedical Research, Cambridge, MA). MGC Human RSPO3 Sequence-Verified cDNA was purchased (GE Dharmacon cat. # MHS6278-202841214). pX330-U6-Chimeric_BB-CBh-hSpCas9 (pX330) was a gift from Feng Zhang (Addgene plasmid # 42230).

pHLsec-HA-hRSPO1-Tev-Fc-Avi-1D4 and pHLsec-HA-hRSPO3-Tev-Fc-Avi-1D4 were constructed through a two-step subcloning strategy. In the first step, human RSPO1 and human RSPO3 were amplified by PCR with forward primers pCX-RSPO1-F (5’-GAG GCT AGC ACC ATG CGG CTT GGG CTG TGT G-3’) or pCX-RSPO3-F (5’-GAG GCT AGC ACC ATG CAC TTG CGA CTG ATT TCT TG-3’), containing an NheI restriction site, and reverse primers pCX-RSPO1-R (5’-TGA GGT ACC AAG GCA GGC CCT GCA GAT GTG-3’) or pCX-RSPO3-R (5’-TGA GGT ACC AAG TGT ACA GTG CTG ACT GAT ACC GA-3’), containing a KpnI restriction site. The products were digested with NheI and KpnI and subcloned into pCX-Tev-Fc digested at the same sites. In the second step, a fragment containing RSPO1 or RSPO3 followed by two tandem Tev cleavage sites, a linker and the Fc domain of human IgG was amplified by PCR from pCX-RSPO1-Tev-Fc or pCX-RSPO3-Tev-Fc, respectively, using forward primers pHL-SEC-RSPO1-F-gibson (5’-CGA CGT GCC CGA CTA CGC CAC CGG TAA CCT GAG CCG GGG GAT CAA GGG G-3’) or pHL-SEC-RSPO3-F-gibson (5’-CGA CGT GCC CGA CTA CGC CAC CGG TAA CCT GCA AAA CGC CTC CCG GG-3’) and reverse primer pHL-SEC-FC-R-gibson-no-KpnI (5’-ACC ACC GGA ACC TCC GGT ACT TTT ACC CGG AGA CAG GGA GA-3’). The forward and reverse primers contained 24 base pair (bp) overhangs complementary to pHLsec-HA-Avi-1D4 upstream of the unique AgeI site and downstream of the unique KpnI site in the vector, respectively. The reverse primer contained a mutation that eliminated the KpnI site in pHLsec-HA-Avi-1D4, hence retaining only one KpnI site between RSPO1 or RSPO3 and the Tev cleavage sites in the resulting construct. The PCR products were subcloned by Gibson assembly (using Gibson Assembly Master Mix, NEB Cat. # E2611L) into pHLsec-HA-Avi-1D4 digested with AgeI and Acc65I (an isoschizomer of KpnI) to produce pHLsec-HA-RSPO1-Tev-Fc-Avi-1D4 and pHLsec-HA-RSPO3-Tev-Fc-Avi-1D4, which contain a single AgeI site upstream and a single KpnI site downstream of the RSPO coding sequence. Henceforth, we refer to the vector backbone of this new constructs as pHLsec-HA-Tev-Fc-Avi-1D4.

Human RSPO1 and RSPO3 mutants and chimeras (Figs. 2a, 3a and Supplementary Data File 2) were generated synthetically as gBlocks Gene Fragments (IDT), flanked at the 5’ and 3’ ends, respectively, by 24 bp overhangs overlapping the sequence upstream of the unique AgeI site and downstream of the unique KpnI site in the HLsec-HA-Tev-Fc-Avi-1D4 vector. The gBlocks were subcloned into pHLsec-HA-Tev-Fc-Avi-1D4, digested with AgeI and KpnI, using the NEBuilder HiFi DNA Assembly Master Mix (NEB Cat. # E2621L).

To remove the dimerizing Fc tag from RSPO3 ΔTSP/BR, a fragment lacking the TSP and BR domains of RSPO3 was amplified by PCR using forward primer pHL-SEC-RSPO3-F-gibson (sequence described above) and reverse primer pHL-SEC-AVI-1D4-RSPO3FU2-R-gibson (5’-AGA CCG GAA CCA CCG GAA CCT CCG GTA CCC ACA ATA CTG ACA CAC TCC ATA GTA TGG TTG T-3’), containing 24 bp overhangs complementary to pHLsec-HA-Avi-1D4 upstream of the unique AgeI site and downstream of the unique KpnI site in the vector, respectively. The PCR product and pHLsec-HA-Avi-1D4 vector were both digested with AgeI and KpnI, and ligated to produce pHLsec-HA-RSPO3ΔTSP/BR-Avi-1D4.

All constructs were sequenced fully and will be deposited in Addgene.

### Analysis of WNT reporter fluorescence

To measure WNT reporter activity in HAP1-7TGP cells or derivatives thereof, ∼24 hrs before treatment cells were seeded in 96-well plates at a density of 1.5x10^4^ per well and grown in 100 μl of complete growth medium (CGM) 2^18^. Cells were treated for 20-24 hrs with the indicated concentrations of WNT3A CM, untagged recombinant human R-Spondin 1, 2, 3 or 4 (R&D Systems Cat. # 4645-RS, 3266-RS, 3500-RS or 4575-RS, respectively), tagged RSPO1-4 proteins (see below) or CHIR-99021 (CT99021) (Selleckchem Cat. # S2924), all diluted in CGM 2. Cells were washed with 100 μl phosphate buffered saline (PBS), harvested in 30 µl of 0.05% trypsin-EDTA solution (Gemini Bio-Products Cat. # 400-150), resuspended in 120 µl of CGM 2, and EGFP fluorescence was measured immediately by FACS on a BD LSRFortessa cell analyzer (BD Biosciences) using a 488 laser and 505LP, 530/30BP filters, or on a BD Accuri RUO Special Order System (BD Biosciences).

For the experiments shown in Figs. 1c, 2b, 2e, 3b, 3e and 4b-d, cells were treated in duplicate or triplicate wells, fluorescence data for 5,000-10,000 singlet-gated cells was collected, and the median EGFP fluorescence for each well and/or the average +/- standard deviation (SD) of the median EGFP fluorescence from each well (as indicated in the figure legends) was used to represent the data. The results from one representative experiment out of at least two conducted on separate days are presented. For the experiments shown in figures 1a-b, 2c-d, 2f-g, 3c-d, 4a, and Extended Data Figs. 1c-d, cells were treated in single wells and fluorescence data for 5,000-10,000 singlet-gated cells was collected. The median EGFP fluorescence and, when compatible with clarity, the standard error of the median (SEM = 1.253 σ /√*n*,where σ = standard deviation and *n* = sample size) from each well was used to represent the data. Dose-response curves were fitted using the nonlinear regression (curve fit) analysis tool in GraphPad Prism 7 using the [agonist] vs. response –variable slope (four parameters) equation with least squares (ordinary) fit option.

### Construction of mutant HAP1-7TGP cell lines by CRISPR/Cas9-mediated genome editing

Oligonucleotides encoding single guide RNAs (sgRNAs) (Supplementary Data File 3) were selected from one of two published libraries^27,28^ and cloned into pX330 according to a published protocol^29^ (original version of “Target Sequence Cloning Protocol” from http://www.genome-engineering.org/crispr/wp-content/uploads/2014/05/CRISPR-Reagent-Description-Rev20140509.pdf).

Clonal HAP1-7TGP cell lines were established by transient transfection with pX330 containing the sgRNA, followed by single cell sorting as described previously^18^. Genotyping was done as described previously^18^ using the primers indicated in Supplementary Data File 3. To generate triple, quadruple and quintuple knock-out (KO) cell lines, a single clonal cell line with the first desired mutation or mutations was used in subsequent rounds of transfection with pX330 containing additional sgRNAs. To facilitate screening of mutant clones by PCR when targeting two genes simultaneously, we sometimes targeted one of the two genes at two different sites within the same exon or in adjacent exons and amplified genomic sequence encompassing both target sites. Mutant clones were readily identified by the altered size of the resulting amplicon, and the precise lesions were confirmed by sequencing the single allele of each gene present in HAP1 cells.

### Production of tagged RSPO proteins by transient transfection of 293T cells and immunoaffinity-purification from conditioned media (see Extended Data Fig. 1a)

∼24 hours before transfection, 14x10^6^ 293T cells were seeded in each of two T-175 flasks for transfection with each construct, each flask containing 30 ml of CGM 1^18^. Once they had reached 60-80% confluency, the cells in each flask were transfected with 1 ml of a transfection mix prepared as follows: 22.3 µg of pHLsec-HA-RSPO-Tev-Fc-Avi-1D4 construct encoding WT or mutant/chimeric RSPO proteins was diluted in 930 µl of serum-free DMEM (GE Healthcare Life Sicences Cat. # SH30081.01) and 70 µl of polyethylenimine (PEI, linear, MW ∼25,000, Polysciences, Inc. Cat. # 23966) were added from a 1 mg/ml stock (prepared in sterile water, stored frozen and equilibrated to 37°C before use). The transfection mix was vortexed briefly, incubated for 15-20 minutes at room temperature (RT) and added to the cells without replacing the growth medium. ∼16 hrs post-transfection, the cells were washed with 30 ml PBS and the medium was replaced with 28 ml of CD 293 medium (Thermo Fisher Scientific Cat. # 11913019) supplemented with 1x L-glutamine solution (stabilized, Gemini Bio-Products Cat. # 400-106), 1x penicillin/streptomycin solution (Gemini Bio-Products Cat. # 400-109) and 2 mM valproic acid (Sigma-Aldrich Cat. # P4543, added from a 0.5 M stock prepared in water and sterilized by filtration through a 0.22 µm filter) to promote protein expression.

∼90 hrs post-transfection, the CM from each of the two flasks, containing secreted, tagged RSPO protein, was centrifuged for 5 min at 400 x g to pellet detached cells. The supernatant was centrifuged for 5 min at 4000 x g and filtered through 0.45 µm filters (Acrodisc syringe filters with Supor membrane, Pall Corporation) to remove particulates, and was reserved on ice.

Rho 1D4 immunoaffinity resin was prepared by coupling Rho 1D4 purified monoclonal antibody (University of British Columbia, https://uilo.ubc.ca/rho-1d4-antibody) to CNBr-activated sepharose 4B (GE Healthcare Life Sciences Cat. # 17-0430-01). Briefly, 1 g of dry CNBr-activated sepharose 4B was dissolved in 50 ml of 1 mM HCl and allowed to swell. The resin was transferred to an Econo-Pac chromatography column (Biorad Cat. # 7321010) and washed by gravity flow with 50 ml of 1 mM HCl, followed by 50 ml of 0.1 M NaHCO_3_, 0.5 M NaCl, pH 8.5. 14 mg of Rho 1D4 antibody were dissolved in 0.1 M NaHCO_3_, 0.5 M NaCl, pH 8.5, and incubated with the resin overnight, rotating at 4°C. The resin was washed with 50 ml of 0.2 M glycine, pH 8.0, and incubated for 2 hrs in the same buffer, rotating at RT. The resin was washed sequentially with 50 ml each of: 0.1 M NaHCO_3_, 0.5 M NaCl, pH 8.5; 0.1 M NaOAc, 0.5 M NaCl, pH 4.0; 0.1 M NaHCO_3_, 0.5 M NaCl, pH 8.5; PBS, 10 mM NaN_3_. The packed resin was resuspended in an equal volume of PBS, 10 mM NaN_3_ to make a ∼50% slurry, aliquoted and stored at 4°C.

300 µl of the ∼50% slurry of Rho 1D4 resin was added to a 50 ml conical tube containing the CM, and the suspension was incubated 10 hrs rocking at 4°C. Following binding and during all subsequent washes, the resin was collected by centrifugation for 5 min at 400 x g in a swinging bucket rotor. The beads were wash three times at RT with 25 ml PBS by resuspending the beads in buffer and mixing by inverting for ∼1 min. Following the third wash the resin was transferred to a 1.5 ml Eppendorf tube and washed three times with 1.4 ml of PBS, 10% glycerol.

Following the last wash, the buffer was aspirated and the resin was resuspended in 150 µl of PBS, 10% glycerol to obtain a ∼50% slurry. Tagged RSPO protein was eluted by adding 3 µl of a 25 mM stock of 1D4 peptide (NH_3_)-T-E-T-S-Q-V-A-P-A-(COOH)) for a final concentration of 250 µM. Elution was carried out by rotating the tube sideways overnight at 4°C. Following centrifugation of the resin, the eluate was recovered and reserved on ice. The resin was resuspended in 150 µl of PBS, 10% glycerol, and 250 µM 1D4 peptide was added. A second round of elution was carried out for 1 hr at RT. Following centrifugation of the resin, the second eluate was recovered and pooled with the first. The final eluate was centrifuged once again to remove residual resin, and the supernatant was aliquoted, frozen in liquid nitrogen and stored at - 80°C.

### Quantification of tagged RSPO proteins by PAGE (see Extended Data Fig. 1b)

4.5 µl and 13.5 µl of the final eluates containing tagged RSPO proteins were diluted with 4x LDS sample buffer (Thermo Fisher Scientific Cat. # NP0007) supplemented with 50 mM *tris*(2-carboxyethyl)phosphine (TCEP), heated for 10 min at 95°C, and loaded alongside Precision Plus Protein molecular weight standards (Bio-Rad Cat. # 1610373) and bovine serum albumin (BSA) standards (Thermo Fisher Scientific Cat. # 23209) for quantification. Proteins were electrophoresed in NuPAGE 4-12% Bis-Tris gels (Thermo Fisher Scientific) using 1X NuPAGE MES SDS running buffer (Thermo Fisher Scientific Cat. # NP0002).

Gels were fixed in 50% methanol, 7% acetic acid for 30 min, rinsed for 1.5 hrs with several changes of water, stained for 2 hrs with GelCode Blue Stain Reagent (based on colloidal coomassie dye G-250, Thermo Fisher Scientific Cat. # 24590), de-stained in water overnight, and imaged using the Li-Cor Odyssey imaging system. Acquisition parameters for coomassie fluorescence (700 nm channel) were set so as to avoid saturated pixels, and bands with intensities within the linear range of fluorescence for the BSA standards were quantified using manual background subtraction.

### Immunoblot analysis of tagged RSPO proteins (see Extended Data Fig. 1b)

50 ng of tagged RSPO proteins were electrophoresed as described above. Proteins were transferred to nitrocellulose membranes in a Criterion Blotter apparatus (Bio-Rad Cat. # 1704071) using 1X NuPAGE transfer buffer (Thermo Fisher Scientific Cat. # NP0006) containing 10% methanol. Membranes were blocked with Odyssey Blocking Buffer (Li-Cor Cat. # 927-40000), incubated overnight at 4°C with purified anti-HA.11 Epitope Tag primary antibody (BioLegend Cat. # 901501, previously Covance cat. # MMS-101P) diluted 1:1,500 in blocking solution (a 1 to 1 mixture of Odyssey Blocking Buffer (Li-Cor Cat. # 927-40000) and TBST (Tris buffered saline (TBS) + 0.1% Tween-20)), washed with TBST, incubated for 1 hr at RT with IRDye 800CW donkey anti-mouse IgG (H+L) (Li-Cor Cat. # 926-32212) diluted 1:10,000 in blocking solution, washed with TBST followed by TBS, and imaged using the Li-Cor Odyssey imaging system.

### Preparation of figures and statistical analysis

Illustrations were prepared using PowerPoint (Microsoft) and Illustrator CS6 (Adobe). Tables and supplementary files were prepared using Excel and Word (Microsoft). Bar graphs, dose-response graphs and circle graphs were prepared using Prism 7 (GraphPad Software) and statistical analysis was performed using the same software. For comparisons between two datasets, significance was determined by unpaired t test; for comparisons between more than two datasets, significance was determined by one-way ANOVA. Significance is indicated as **** (*p* < 0.0001), ** (*p* < 0.01), * (*p* < 0.05) or ns (not significant). Pictures of gels and immunoblots were only adjusted for contrast and brightness when necessary for clarity using Photoshop CS6 (Adobe), and were arranged in Illustrator CS6.

### Data availability

All data generated or analyzed during this study are included in this published article (and its supplementary information files).

## Acknowledgments

We thank Jan Carette and Rohatgi lab members for input on the project, Henry Ho for pCX-Tev-Fc, Christian Siebold for pHLsec-HA-Avi-1D4 and Feng Cong for RSPO1-GFP. This work was funded by NIH grants DP2 GM105448 and R35 GM118082 to R.R. A.M.L. was supported by the Stanford Dean’s Postdoctoral Fellowship, the Stanford Cancer Biology Program Training Grant and the Novartis sponsored Fellowship from the Helen Hay Whitney Foundation.

## Author Contributions

A.M.L and R.R. conceived the study, designed experiments and analyzed the data. A.M.L. conducted all experiments. A.M.L. and R.R. wrote the manuscript.

## Author Information

The authors declare no conflicting financial interests.

## Supplementary Information

Supplementary Data File 1. List of clonal cell lines used in this study.

Clonal cell lines in which a single or multiple genes were targeted using CRISPR/Cas9 are described in two separate spreadsheets labeled accordingly. For cell lines engineered using CRISPR/Cas9, when more than one clone was generated using the same CRISPR guide, the “Clone Name” column indicates the generic name used throughout the manuscript to describe the genotype, and the “Clone #” column identifies the specific allele in each individual clone. The figures in which each clone was used are also indicated. The “CRISPR guide” column indicates the name of the guide used, which is the same as that of the oligos encoding sgRNAs (see Methods and Supplementary Data File 3). The “Genomic Sequence” column shows 80 bases of genomic sequence (5’ relative to the gene is to the left) surrounding the target site. For each group of clones made using the same CRISPR guide (separated by gray spacers), the “Genomic Sequence” column is headlined by the reference WT genomic sequence (obtained from RefSeq), with the guide sequence colored blue. The site of the double strand cut made by Cas9 is between the two underlined bases. Sequencing results for individual clones are indicated below the reference sequence. Some WT clones are indicated as such and were used as controls. For mutant clones, mutated bases are colored red (dashes represent deleted bases, three dots are used to indicate that a deletion continues beyond the 80 bases of sequence shown, and large insertions are indicated in brackets), and the nature of the mutation, the resulting genotype and any pertinent observations are also described. For clones in which multiple genes were targeted, the CRISPR guide or pair of guides used (in some cases two different guides were used simultaneously to target adjacent sites in the same gene), genomic sequence, mutation, genotype and observations pertaining to each of the targeted genes are designated “1”, “2”, “3” and so on in the column headings, and are shown under spacers of different colors, respectively.

Supplementary Data File 2. Nucleotide sequences of RSPO1 and RSPO3 WT, mutant and chimeric constructs used in this study.

Lowercase sequences overlap the vector sequence upstream of the unique AgeI site and downstream of the unique KpnI site in the pHLsec-HA-Tev-Fc-Avi-1D4 vector. Uppercase sequences encode RSPO1 or RSPO3. For point mutants, mutated codons relative to WT are underlined.

Supplementary Data File 3. List of oligonucleotides and primers used for generation and characterization of clonal cell lines engineered using CRISPR/Cas9.

The names and sequences of pairs of oligonucleotides encoding sgRNAs (which were cloned into pX330) are shown in the first and second columns, respectively. The names and sequences of pairs of primers used to amplify corresponding genomic regions flanking sgRNA target sites are shown in the third and fourth columns, respectively. The names and sequences of single primers used for sequencing of the amplified target sites are shown in the fifth and sixth columns, respectively.

**Extended Data Figure 1.**
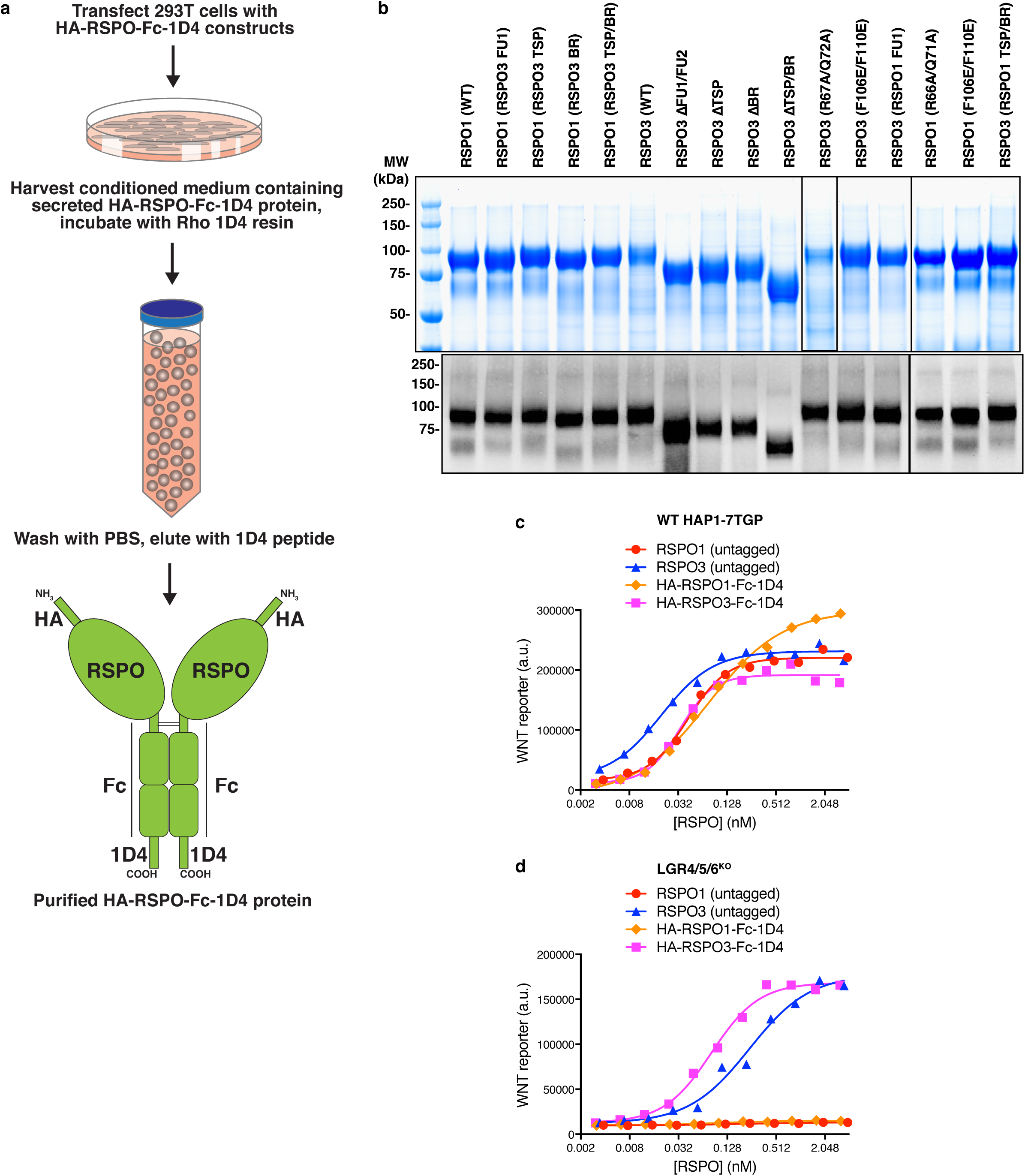
Affinity purification and functional characterization of recombinant RSPO proteins used in this study. a. Summary of a new experimental strategy for the rapid, one-step purification of secreted WT and mutant RSPO proteins containing an HA epitope tag at the N-terminus and a dual Fc-1D4 tag at the C-terminus. The Fc fusion stabilized the various RSPO mutants used in the study, the 1D4 tag enabled immunoaffinity purification under native conditions, and the HA tag allowed quantitative immunoblotting to determine relative ligand concentrations and ensure that each RSPO ligand was produced as a full-length species. See Methods for details. b. Equal volumes (13.5 μl each) of the final eluate for each purified RSPO protein were resolved by polyacrylamide gel electrophoresis (PAGE) and stained with coomassie (top panel). Proteins were quantified by fluorimetry using the Licor Odyssey scanner and then equal mass amounts of each protein were analyzed by immunoblotting against the HA tag (bottom panels). c and d. Dose-response curves comparing untagged RSPOs to RSPOs tagged with HA and Fc-1D4 tags (shown in b) in WT HAP1-7TGP (c) and LGR4/5/6^KO^ (d) cells in the presence of 1.43% WNT3A CM. Each symbol represents the median WNT reporter fluorescence from 5,000 cells.

**Extended Data Table 1.** Relative gene expression level in HAP1 cells of selected genes discussed in this work. RPKM (<u>R</u>eads <u>P</u>er <u>K</u>ilobase of transcript per <u>M</u>illion mapped reads) values from duplicate RNAseq datasets generated as described previously^18^ from two different passages of WT HAP1 cells are shown. Groups of paralogues or genes with redundant function are shaded in alternating colors to facilitate comparisons.

| Gene | RPKM |  |  |
| --- | --- | --- | --- |
|  | Replicate 1 | Replicate 2 | Average |
| *LGR4* | 160.61 | 174.69 | 167.65 |
| *LGR5* | 0.02 | 0.00 | 0.01 |
| *LGR6* | 0.02 | 0.00 | 0.01 |
| *ZNRF3* | 30.9 | 33.3 | 32.1 |
| *RNF43* | 0.12 | 0.08 | 0.1 |
| *GPC1* | 49.55 | 47.53 | 48.54 |
| *GPC2* | 4.17 | 4.79 | 4.48 |
| *GPC3* | 170.22 | 144.37 | 157.29 |
| *GPC4* | 209.39 | 229.86 | 219.63 |
| *GPC5* | 0.1 | 0.1 | 0.1 |
| *GPC6* | 13.88 | 14.90 | 14.39 |
| *SDC1* | 51.37 | 47.88 | 49.63 |
| *SDC2* | 11.42 | 9.2 | 10.31 |
| *SDC3* | 43.58 | 50.64 | 47.11 |
| *SDC4* | 8.16 | 8.21 | 8.18 |

